# Stable m^7^G Cap-Distal 5′UTR Hairpin Structure Mediates Distinct 40S and 60S Binding Dynamics

**DOI:** 10.1101/2021.12.29.474417

**Authors:** Hongyun Wang, Anthony Gaba, Xiaohui Qu

## Abstract

The 5′ untranslated region (UTR) of diverse mRNAs contains secondary structures that can influence protein synthesis by modulating the initiation step of translation. Studies support the ability of these structures to inhibit 40S subunit recruitment and scanning, but the dynamics of ribosomal subunit interactions with mRNA remain poorly understood. Here, we developed a reconstituted *Saccharomyces cerevisiae* cell-free translation system with fluorescently labeled ribosomal subunits. We applied this extract and single-molecule fluorescence microscopy to monitor, in real time, individual 40S and 60S interactions with mRNAs containing 5’ UTR hairpin structures with varying thermostability. In comparison to mRNAs containing no or weak 5′UTR hairpins (ΔG >= -5.4 kcal/mol), mRNAs with stable hairpins (ΔG <= -16.5 kcal/mol) showed reduced numbers of 60S recruitment to mRNA, consistent with the expectation of reduced translation efficiency for such mRNAs. Interestingly, such mRNAs showed increased numbers of 40S recruitment events to individual mRNAs but with shortened duration on mRNA. Correlation analysis showed that these unstable 40S binding events were nonproductive for 60S recruitment. Furthermore, although the mRNA sequence is long enough to accommodate multiple 40S, individual mRNAs are predominantly observed to engage with a single 40S at a time, indicating the sequestering of mRNA 5’ end by initiating 40S. Altogether, these observations suggest that stable cap-distal hairpins in 5’ UTR reduce initiation and translation efficiency by destabilizing 40S-mRNA interactions and promoting 40S dissociation from mRNA. The premature 40S dissociation frees mRNA 5′-end accessibility for new initiation events, but the increased rate of 40S recruitment is insufficient to compensate for the reduction of initiation efficiency due to premature 40S dissociation. This study provides the first single-molecule kinetic characterization of 40S/60S interactions with mRNA during cap-dependent initiation and the modulation of such interactions by cap-distal 5’ UTR hairpin structures.

## INTRODUCTION

The canonical cap-dependent translation initiation pathway in eukaryotic cells occurs through a scanning mechanism that begins with 40S ribosomal subunit association with a ternary complex (eukaryotic initiation factor (eIF) 2, GTP, and methionyl initiator tRNA) and the eIFs 1, 1A, 3, and 5, which collectively forms a 43S pre-initiation complex (PIC). The eIF4F complex (eIF4E/4G/4A) binds to the 5′-terminal 7-methylguanosine (m^7^G) cap structure on mRNA and facilitates 43S recruitment to the mRNA 5′ end. Then the 43S scans the mRNA 5′-untranslated region (UTR) in a 5′ to 3′ direction. 43S recognition of a start codon stops the scanning 43S and leads to release of eIFs, PIC structural rearrangements, and 60S subunit joining to form the 80S ribosome, which then proceeds to the peptide elongation stage (1). 43S scanning is thought to be modulated by 5′UTR secondary structures, which are abundant in naturally occurring eukaryotic mRNAs and can exist in diverse forms such as hairpins and G-quadruplex structures (2-5). The 5′UTRs of mRNAs that encode proteins in cancer-associated and cell growth regulation pathways are enriched with secondary structures (6-9), underscoring the importance of understanding effects of 5′UTR-localized secondary structures on dynamics of ribosome-mRNA interactions.

Biochemical studies demonstrated that 5′UTR hairpin structure could regulate translation in a manner that was dependent on both hairpin position and strength (10). Hairpins with moderate stabilities (−10–30 kcal/mol) and positioned within 12 nt from the m^7^G cap inhibited translation in cell-free translation reactions (10) and in cells (11). While moderately stable -30 kcal/mol hairpins positioned further (50–60 nt) from the m^7^G cap did not inhibit translation in cell extract (10) or in COS cells (12), a hairpin 72 nt from the m^7^G cap did inhibit translation when its stability was increased to -61 kcal/mol (10). Recruitment of the 40S subunit to mRNA, however, was not inhibited by the -61 kcal/mol hairpin but was inhibited with a -30 kcal/mol hairpin positioned 12 nt from the m^7^G cap in cell-free translation reactions (10) and with a -7.4 kcal/mol hairpin positioned 10 nt from the m^7^G cap in a reconstituted translation initiation system with purified translation initiation factors (13). These studies have suggested that moderately stable hairpins can inhibit 43S binding to mRNA when positioned proximal to the m^7^G cap, whereas hairpins distal to the m^7^G cap do not inhibit 43S recruitment but can inhibit translation by impeding 43S scanning. The dynamics of ribosomal subunit interactions with mRNA, however, remain poorly understood because the intrinsic stochasticity of individual initiation events challenge traditional biochemical approaches.

To overcome these challenges, we developed a reconstituted eukaryotic cell-free translation system with fluorescently labeled ribosomal subunits. By combining the reconstituted extract, well-designed reporter mRNAs, and single-molecule fluorescence microscopy, we were able to establish a single-molecule assay that allowed kinetic observation of individual 40S and 60S interactions with mRNA during active cap-dependent translation. This single-molecule assay preserves the translation characteristics of bulk assays, including the capability of polypeptide synthesis, the m^7^G cap stimulation of translation, and the dependence of 60S binding to mRNAs on the presence of a start codon and 40S subunit. When applying this assay to reporter mRNAs containing cap-distal 5’ UTR hairpin structures with varying thermostability, we observed that the different hairpin stabilities caused distinct 40S and 60S binding dynamics. Our single-molecule observation of ribosome recruitment dynamics suggests that stable cap-distal hairpins down-regulate 60S recruitment by destabilizing 40S subunit interactions with mRNA and promoting premature 40S dissociation. This assay can be easily expanded to different eukaryotic translation systems and should enable a wide range of applications for studying the regulation of ribosome recruitment dynamics by the large variety of regulatory elements that have been identified.

## RESULTS

### A Single-Molecule Method to Monitor 40S and 60S Ribosomal Subunit Interactions with mRNA

Mechanisms and kinetics of ribosome recruitment to mRNA during cap-dependent initiation remain incompletely understood because traditional methods cannot resolve individual ribosome-mRNA binding events, which are intrinsically stochastic and potentially heterogeneous. To investigate these dynamics, new methods are needed to resolve the individual asynchronous ribosome interactions with mRNA during cap-dependent translation. Here, we developed a *Saccharomyces cerevisiae* (*S. cerevisiae*) cell-free translation system reconstituted with fluorescently labeled and purified 40S (LD550-40S) and 60S (LD650-60S) ribosomal subunits, ribosome depleted extract (RDE), and extract factors (EFs) (Figure 1A; Materials and Methods). The 40S and 60S ribosomal subunits were labeled with triplet-state quencher-conjugated maleimide dyes, LD550 and LD650, respectively (14). When comparing the fluorescence from equal molar amounts of labeled ribosomal particles versus the free maleimide dyes, fluorescence intensity levels were approximately two-fold greater from LD550-40S and LD650-60S compared to the maleimide dyes alone (Supplementary Fiigure 1), indicating an average labeling ratio of two fluorophores per individual 40S and 60S. Analysis of 18S and 25S rRNAs in the reconstituted extract components confirmed that the fluorescently labeled ribosomal particles (LD550-40S and LD650-60S) were intact and that the RDE and EFs were free of ribosomal particles (Figure 1B).

**Figure 1.**
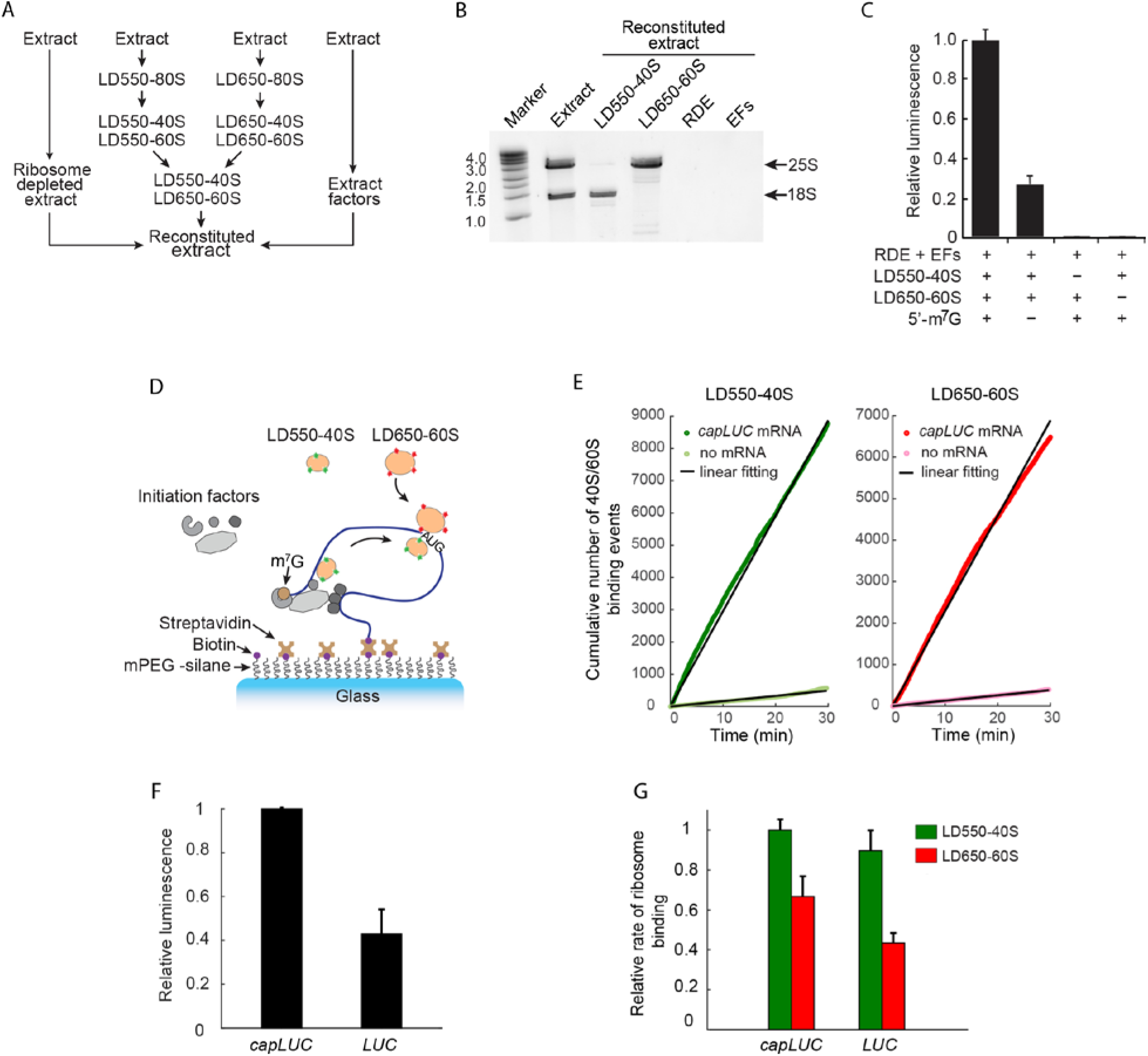
A reconstituted cell-free translation system with fluorescently labeled ribosomal subunits. (*A*) Flow chart for reconstitution of *S. cerevisiae* extract with fluorescently labeled 40S and 60S. (*B*) Analysis of 18S and 25S rRNAs in reconstituted extract components. Total RNA from extract and the reconstituted extract components LD550-40S, LD650-60S, ribosome-depleted extract (RDE), and extract factors (EFs) was analyzed on a denaturing gel. Amounts loaded are equivalent to those in a 10 μl translation reaction with reconstituted extract and 0.1μl of starting extract. 18S and 25S rRNAs are indicated, as are the sizes of an RNA ladder (marker). (*C*) Relative levels of firefly luciferase production in bulk cell-free translation reactions with reconstituted extract components. All luminescence readings were normalized to the activity of an internal control Renilla luciferase (materials and methods). Mean values and standard deviations from two independent translation reactions are given. (*D*) Schematic of the single-molecule assay. 3’-end biotinylated mRNA molecules are end-tethered to a PEGylated detection surface via biotin-streptavidin interactions. Reconstituted extract is flowed into the detection chamber and the dynamics of LD550-40S and LD650-60S interactions with single mRNA molecules are tracked. (*E*) Representative results for the cumulative number of LD550-40S (green) and LD650-60S (red) binding events per imaging area over time with *capLUC* mRNA translation (dark color) or without mRNA (light color). 40S and 60S binding with and without mRNA represent specific binding to mRNA and nonspecific binding to single-molecule detection surface, respectively. The slope of linear fitting to each cumulative plot yielded the average rate of 40S/60S binding per imaging area as the following: 4.93±0.01 (for specific LD550-40S binding), 0.25±0.01 (for nonspecific LD550-40S binding), 3.83±0.01 (for specific LD650-60S binding), 0.21±0.01 (for nonspecific LD650-60S binding). (*F*) Relative levels of firefly luciferase production in single-molecule chambers with *capLUC* or *LUC* mRNA. Luminescence readings were normalized to the activity in chambers with *capLUC* mRNA, which was arbitrarily set to 1.0. (*G*) Relative rate of LD550-40S and LD650-60S binding per imaging area in single-molecule chambers with *capLUC* or *LUC* mRNA. The binding rates were normalized to the binding rate of LD550-40S in chambers with *capLUC* mRNA, which was arbitrarily set to 1.0.

Bulk *in vitro* translation using the reconstituted extract and firefly luciferase-encoding reporter mRNAs showed luciferase activity levels that were ∼3.5-fold greater with the m^7^G-capped mRNA (*capLUC*) compared to uncapped *LUC* mRNA (Figure 1C). This magnitude of cap-stimulated translation activity is consistent with previous observations with bulk cell-free translation reactions using micrococcal nuclease-treated *S. cerevisiae* extract (15). Omission of either ribosomal subunit in the reconstituted extract reduced the luminescence reading of luciferase activity to the background level (Figure 1C), indicating that addition of both subunits was necessary for *capLUC* mRNA translation. These results indicate that the reconstituted extract with fluorescently labeled LD550-40S and LD650-60S enabled protein synthesis, produced enzymatically active proteins, and preserved translation through a cap-dependent pathway.

To facilitate single-molecule imaging, 3′-end biotinylated mRNA was immobilized to a streptavidin-coated detection surface in a flow channel (Figure 1D). In channels filled with reconstituted extract, the dynamics of 40S and 60S binding with individual immobilized mRNAs are detected using total internal reflection fluorescence (TIRF) imaging (Figure 1D). We first tested the single-molecule assay for translation activity by using *capLUC* mRNA and measuring the cumulative number of LD550-40S and LD650-60S binding events per imaging area over time (Figure 1E). The rate of 40S or 60S binding per imaging area was determined as the slope yielded from a linear fit to each cumulative curve. We observed that the rates of LD550-40S and LD650-60S specific binding to *capLUC* mRNA were approximately 19-fold faster than the rates of their nonspecific binding to a detection surface lacking mRNA (Figure 1E). These results indicate that the levels of specific LD550-40S and LD650-60S binding to 3′-end-immobilized *capLUC* mRNA were significantly above the levels of nonspecific ribosome binding to detection surface.

Following single-molecule detection with *capLUC* mRNA, the translation mix can be collected from single-molecule channels and assayed in bulk for luciferase activity to assess the amount of luciferase proteins synthesized during single-molecule detection. These translation mixes showed luminescence readings that were ∼2.5-fold greater with *capLUC* mRNA than with uncapped *LUC* mRNA (Figure 1F). Thus, the extent of cap-stimulation was similar with translation in single-molecule (Figure 1F) and bulk (Figure 1C) conditions.

Given the observed ∼2.5-fold cap stimulation of single-molecule translation (Figure 1F), we expected a similar extent of cap-stimulated LD550-40S and LD650-60S recruitment to *capLUC* mRNA if the ribosomal subunit recruitment levels linearly correlated with luciferase production. However, the m^7^G cap stimulated the rates of LD550-40S and LD650-60S recruitment to *capLUC* mRNA by only ∼1.1-fold and ∼1.5-fold, respectively (Figure 1G). These results are consistent with previous bulk observations indicating that 40S recruitment to uncapped mRNA involves a nonproductive pathway (13) and with previous ribosome profiling studies indicating that extensive 80S initiation occurs at alternative AUG and non-AUG (near-cognate) start codons (16-18). There are >100 AUG codons and near-cognate start codons in the ∼1.9 kb *LUC* mRNA sequence. Due to the vast amounts of potential alternative initiation sites in the *LUC* sequence, *capLUC* mRNA was not an optimal substrate for single-molecule analysis of cap-dependent ribosome-mRNA interactions.

### LD650-60S recruitment to *capCAA*_*AUG*_ mRNA occurs through the canonical initiation pathway

To facilitate better mechanistic understanding, we designed mRNA sequences that lack alternative AUG and near-cognate initiation sites. Specifically, we generated synthetic 119 nt capped CAA repeat mRNAs with either a single AUG start codon positioned 60–62 nt downstream of the mRNA 5′-end (*capCAA*_*AUG*_) or with the start codon mutated to AAC (*capCAA*_*AAC*_) (Figure 2A). CAA repeat mRNA sequences do not contain alternative AUG or near cognate start codons, are predicted to lack secondary structures (19, 20), and have been useful in previous bulk kinetic studies of RNA helicase function (21).

**Figure 2.**
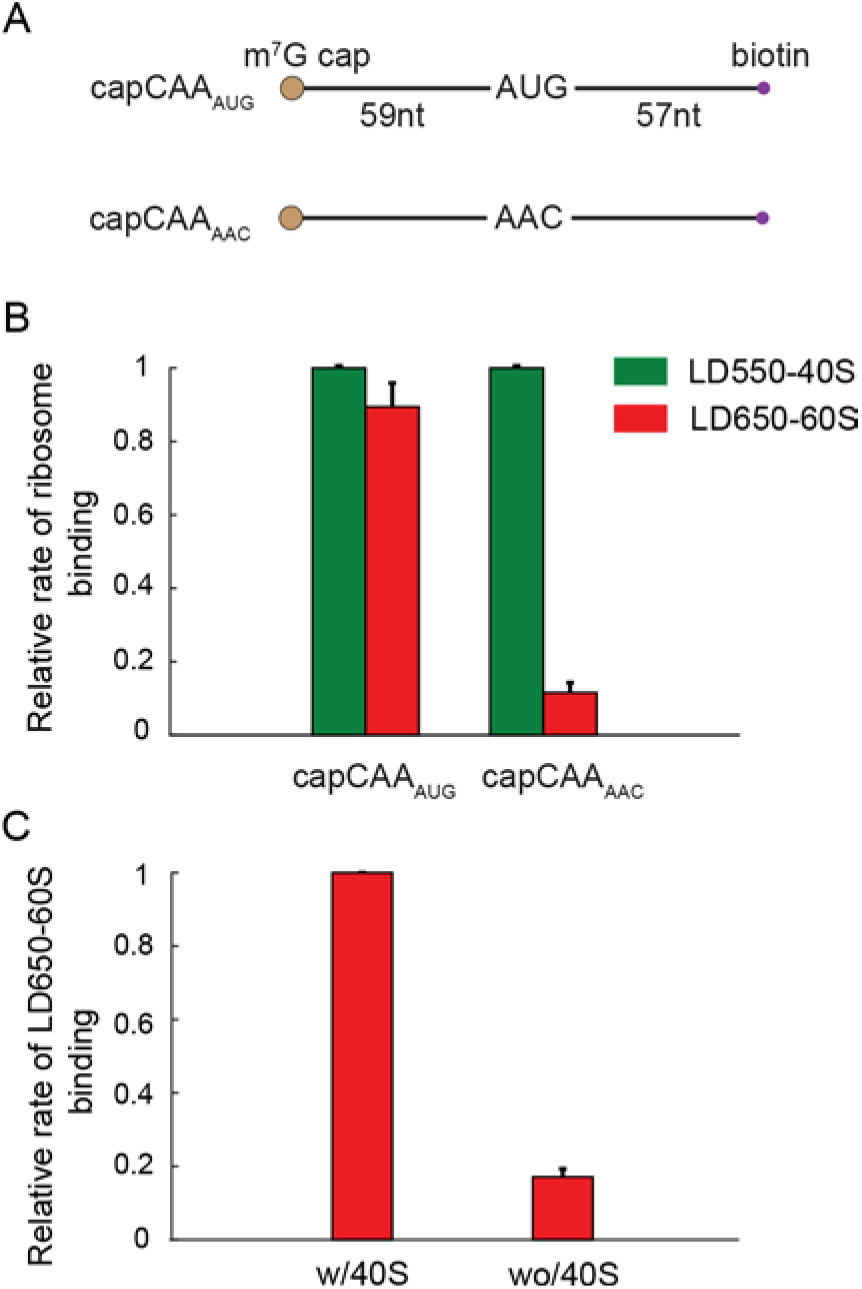
LD650-60S binding to *capCAA*_*AUG*_ mRNA is AUG codon- and LD550-40S-dependent. (*A*) Schematic representation of *capCAA*_*AUG*_ and *capCAA*_*AAC*_ mRNAs. (*B*) Relative rate of LD550-40S and LD650-60S binding per imaging area to *capCAA*_*AUG*_ vs. *capCAA*_*AAC*_ mRNAs. The LD650-60S binding rates (red) are normalized to the LD550-40S binding rates (green), which are arbitrarily set to 1.0 in each condition. (*C*) Relative rate of LD650-60S binding per imaging area to *capCAA*_*AUG*_ mRNA in reactions with full translation extract vs. LD550-40S-lacking extract. The LD650-60S binding rate in the absence of LD550-40S (‘wo/40S’) is normalized to the rate for the full extract (‘w/40S’), which is arbitrarily set to 1.0.

40S and 60S recruitment to CAA repeat mRNA was examined for features of cap-dependent initiation. We first compared 60S binding kinetics to *capCAA*_*AUG*_ and *capCAA*_*AAC*_ mRNAs. The rate of LD650-60S recruitment to the AUG-lacking *capCAA*_*AAC*_ mRNA was reduced ∼8-fold compared to the AUG-containing *capCAA*_*AUG*_ mRNA (Figure 2B), indicating the strong dependence of 60S recruitment on the presence of AUG start codon. We next tested for the dependence of 60S recruitment on 40S recruitment by comparing the kinetics of 60S binding to *capCAA*_*AUG*_ mRNA in reactions with either the complete reconstituted extract or the extract that lacked LD550-40S. The 60S recruitment rates were ∼6-fold slower in reactions with 40S-lacking extract compared to extract with both 40S and 60S (Figure 2C). Collectively, these results indicate that LD650-60S recruitment to *capCAA*_*AUG*_ mRNA was strongly dependent on both the presence of the AUG start codon and 40S recruitment to mRNA. Therefore, 60S interactions with *capCAA*_*AUG*_ mRNA appeared to be dominated by the canonical initiation pathway, wherein 60S is recruited to 40S at the start codon.

### Stable 5’ end-distal hairpins reduce LD650-60S recruitment to *hp-capCAA*_*AUG*_ mRNA

5’ end-distal secondary structures in mRNA are known to reduce translation efficiency (10, 11, 22). However, the relationship between mRNA secondary structure and ribosomal subunit recruitment dynamics remains poorly understood. To investigate these subunit binding dynamics, we designed hairpin-containing *capCAA*_*AUG*_ mRNAs (*hp-capCAA*_*AUG*_) containing a 5′ end-distal hairpin positioned 47 nt downstream of the 5′-m^7^G cap and 12 nt upstream of the AUG start codon (Figure 3A). The hairpins had a common 4 nt ACAC loop region and a stem region containing either 4, 8, 12, or 16 GC base pairs with thermostabilities ranging from -5.4 kcal/mol to -41.3 kcal/mol (20) (Supplementary Tables 1 and 2). The *capCAA*_*AUG*_ mRNA will also be referred to as the 0 GC *hp-capCAA*_*AUG*_ mRNA hereinafter.

**Figure 3.**
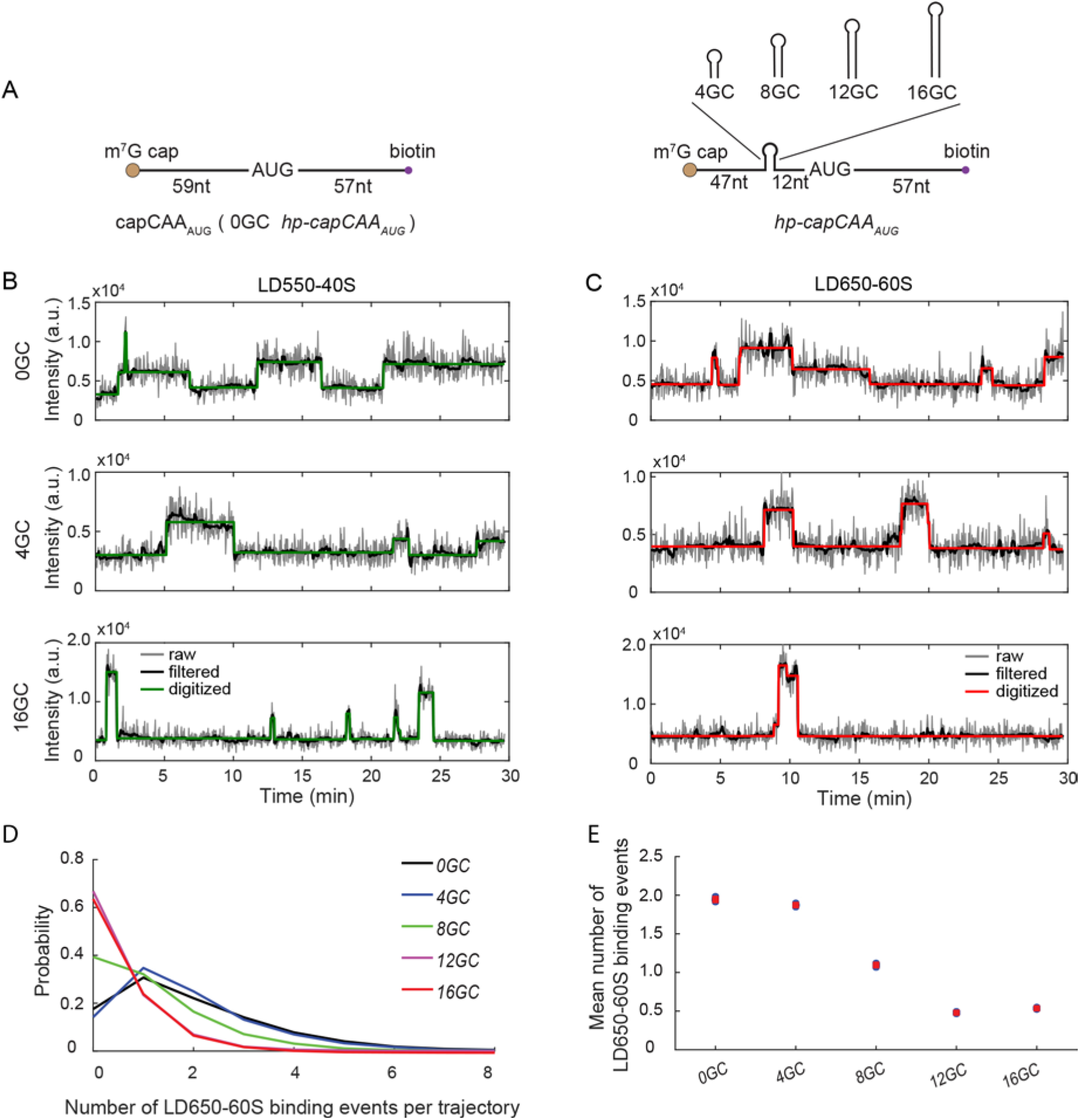
Cap-distal hairpin structures in 5′ UTR reduce LD650-60S binding to single *hp-capCAA*_*AUG*_ mRNAs. (*A*) Schematic representation of *hp-capCAA*_*AUG*_ mRNAs. The numbers of GC base pairs in the hairpins of *hp-capCAA*_*AUG*_ mRNAs are indicated. *capCAA*_*AUG*_ mRNA is also referred to as 0GC *hp-capCAA*_*AUG*_ mRNA hereinafter. (*B* and *C*) Representative trajectories for LD550-40S (*B*) and LD650-60S (*C*) ribosomal subunit binding dynamics on single *hp-capCAA*_*AUG*_ mRNAs. Raw, filtered, and digitized data are shown as indicated (Materials and Methods). (*D* and *E*) Probability distributions (*D*) and mean values (*E*) of LD650-60S binding number per trajectory. (Mean ± SEM: 1.94±0.02 (0GC), 1.87±0.01 (4GC), 1.10±0.01 (8GC), 0.48±0.01 (12GC), 0.54±0.01 (16GC)) Numbers of trajectories analyzed: 0 GC, n = 7573; 4 GC, n = 13310; 8 GC, n = 9582; 12 GC, n = 8699; 16 GC, n = 12040.

In representative single molecule trajectories of LD550-40S (Figure 3B) and LD650-60S (Figure 3C) binding to *hp-capCAA*_*AUG*_ mRNAs, each instantaneous fluorescence increase and decrease represent subunit binding and dissociation, respectively, to single mRNA molecules. The measured numbers of LD650-60S binding events per trajectory decreased with increasing hairpin stability (Figures 3 D,E), indicating that the hairpins negatively regulated LD650-60S recruitment to the *hp-capCAA*_*AUG*_ mRNAs. This observation is consistent with the ability of 5’UTR secondary structures to inhibit translation efficiency, presumably by inhibiting 40S scanning (10, 11, 13).

### Stable 5’ end-distal hairpins promote premature LD550-40S dissociation from mRNA

We analyzed the kinetics of 40S recruitment to *hp-capCAA*_*AUG*_ mRNAs to investigate the modulation of 40S interactions with mRNA by 5’ end-distal hairpins. Surprisingly, we found that the numbers of LD550-40S binding events per trajectory with the 8–16 GC *hp-capCAA*_*AUG*_ mRNAs were 2-fold greater than with the 0GC and 4GC mRNAs (Figures 4 A,B). The increased 40S recruitment to 8–16 GC *hp-capCAA*_*AUG*_ mRNAs appeared contradictory to the decreased 60S recruitment to these mRNAs (Figures 3 D,E). To further examine the LD550-40S binding events, dwell times of individual LD550-40S binding events were measured. The cumulative dwell time histograms showed two types of 40S binding events, manifested by the biphasic nature of the histograms (Figure 4C). Specifically, there was a short-lived mode lasting less than ∼100 sec and a long-lived mode lasting longer than ∼100 sec (Figure 4C). 40S binding to the 0-4 GC *hp-capCAA*_*AUG*_ mRNAs was dominated by the long-lived type, while 40S binding to the 8-12 GC *hp-capCAA*_*AUG*_ mRNAs was dominated by the short-lived type (Figure 4C). Overall, the mean dwell time of 40S binding events decreased by approximately 5-fold in presence of the stable 8–16 GC base pair hairpins (Figure 4D). To examine the relation between the two types of 40S binding events and LD650-60S recruitment, heatmaps were generated to plot the numbers of LD650-60S binding events (y-axis) as a function of individual LD550-40S binding event dwell times (x-axis) (Figure 5). Clearly, only the long-lived 40S binding events are productive for 60S recruitment. Collectively, these results demonstrated that there are two types of 40S binding events, among which only the long-lived ones (>100 sec) are productive for 60S recruitment. Stable 5’ cap-distal hairpins promoted premature 40S dissociation and led to the significantly reduced level of the long-lived and productive 40S binding events.

**Figure 4.**
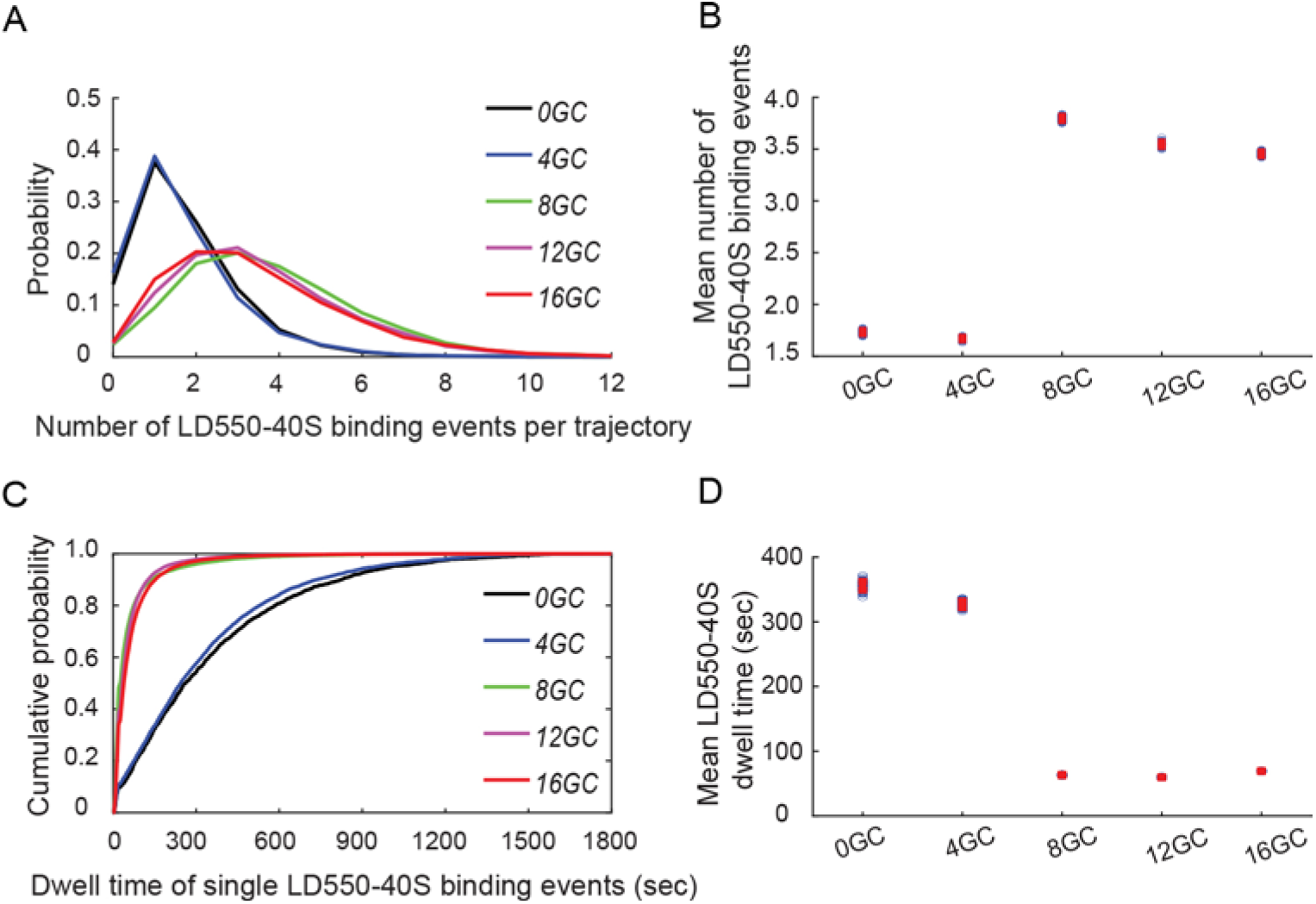
The 8–16 GC base pair 5′ UTR hairpins modulate LD550-40S binding dynamics. (*A* and *B*) Probability distributions (*A*) and mean values (*B*) of LD550-40S binding number per trajectory with *hp-capCAA*_*AUG*_ mRNAs. (Mean ± SEM: 1.73±0.02 (0GC), 1.67± 0.01 (4GC), 3.80±0.02 (8GC), 3.55±0.02 (12GC), 3.45±0.02 (16GC)). Numbers of trajectories analyzed are the same as in Figures 3D and 3E. (*C* and *D*) Cumulative probability distributions (*C*) and mean values (*D*) of LD550-40S single binding event dwell times. Mean dwell time (sec): 356±7 (0GC), 327±5 (4GC), 63±1 (8GC), 60±1 (12GC), 69±1 (16GC). Numbers of LD550-40S single binding events analyzed: 0 GC, n = 1776; 4 GC, n = 5084; 8 GC, n = 16274; 12 GC, n = 18073; 16 GC, n = 22171.

**Figure 5.**
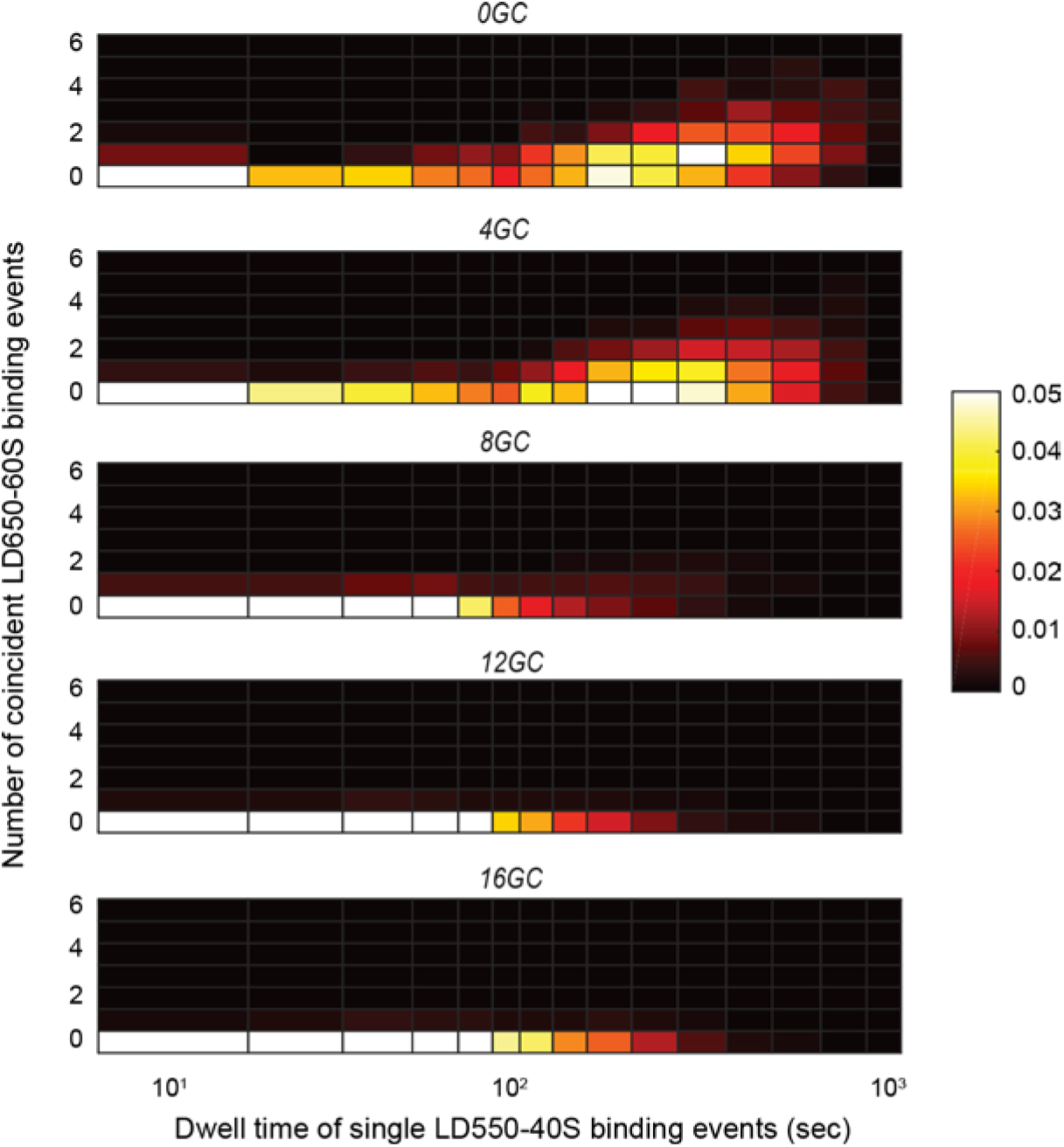
Only stable 40S binding events are correlated with 60S recruitment. Heatmaps represent the correlation between the dwell times of individual LD550-40S binding events (x-axis) and the numbers of recruited LD650-60S binding events (y-axis). The color-gradient represents the probability of observing y numbers of LD650-60S binding events for LD550-40S binding events with dwell times of x seconds. Numbers of LD550-40S single binding events analyzed are the same as in Figure 4.

### LD550-40S binding kinetics suggest mRNA 5’ end sequestration by scanning 40S

Alternative 40S scanning models involve distinct durations of 43S/eIF4F association with 5′ cap (23). In the cap-severed model, the 43S/eIF4F interactions with 5’ cap are disrupted when scanning begins. In the cap-tethered model, 43S/eIF4F interactions with 5’ cap are preserved until initiation terminates at the AUG start codon, limiting mRNAs to one on-going initiation event at a time. To examine CAA repeat mRNAs for these modes of initiation, we measured the numbers of LD550-40S that simultaneously occupied individual mRNAs. Translating 80S ribosomes with open or occupied ribosomal A sites generate ribosome protected footprint sizes of 20–22 nt or 27–29 nt, respectively (24). CAA-repeat mRNAs contain 59 nt upstream and 57 nt downstream from the AUG start codon (Figure 2A). Therefore, CAA-repeat mRNAs should be long enough to accommodate at least 1-2 scanning 40S and 1-2 translating 80S simultaneously. If CAA-repeat mRNA translation followed the cap-severed model, the single-molecule trajectories would be dominated by events of two or more 40S co-existing on the same mRNA, either in the form of the 43S scanning particle or the translating 80S. However, the probability distributions of the numbers of simultaneously bound LD550-40S per mRNA showed that the majority (∼75%) of 40S binding events on individual mRNAs were with a single 40S (Figure 6). This observation clearly showed that CAA-repeat mRNA translation did not follow the cap-severed model. Together with the 40S binding dwell time analysis (Figure 4D), our data showed that 43S/eIF4F interactions with 5’ cap can persist for at least ∼60–350 sec on *hp-capCAA*_*AUG*_ mRNAs, leading to mRNA 5’ end sequestration in the same time scale. Although CAA-repeat mRNA translation is more in line with the cap-tethered model, our data cannot test whether the disruption of the interactions between 43S/eIF4F and 5’ cap is coincidental with initiation termination. Our data cannot rule out he possibilities that such interactions may be disrupted prior to initiation termination or persist into the elongation stage.

**Figure 6.**
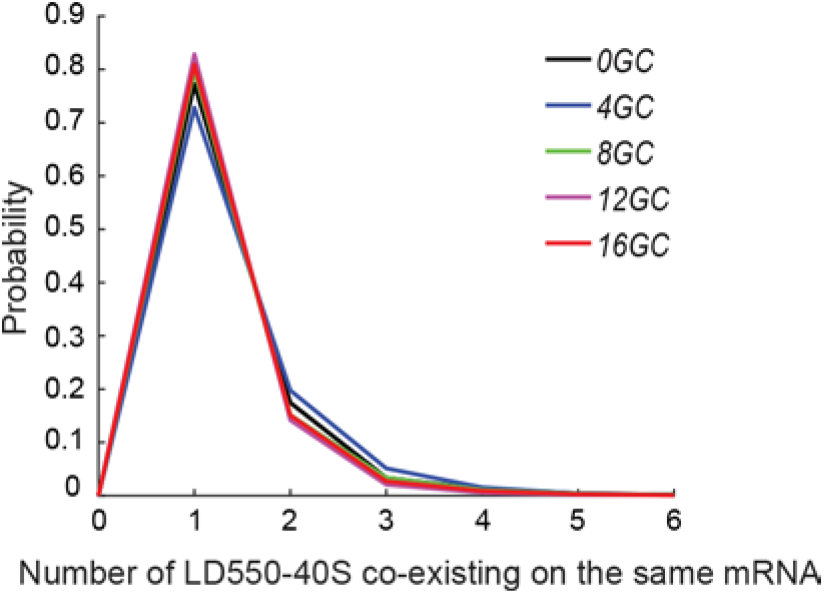
Probability distributions of the numbers of simultaneously bound LD550-40S per *hp-capCAA*_*AUG*_ mRNA. Numbers of LD550-40S binding clusters analyzed: 0 GC, n = 2299; 4 GC, n = 6980; 8 GC, n = 20399; 12 GC, n = 21815; 16 GC, n = 27406.

## DISCUSSION

The mechanisms and dynamics of ribosome interactions with mRNA during cap-dependent initiation remain poorly understood. To monitor the dynamics of 40S and 60S interactions with mRNA, we developed a cell extract fractionation method that enables extract reconstitution with purified and fluorescently labeled ribosomal subunits. By measuring luciferase activity in translation reactions with reconstituted *S. cerevisiae* extract, we showed m^7^G cap-dependent luciferase synthesis by fluorescently labeled ribosomes in bulk and single-molecule conditions (Figures 1*C* and 1*F*). These results, and observations of strong AUG start codon- and 40S-dependent 60S recruitment to *capCAA*_*AUG*_ mRNA (Figures 2B and 2C), indicated that initiation in the reconstituted system occurred through an m^7^G cap-dependent scanning mechanism. These features of the reconstituted system allowed us to investigate the dynamics of ribosome recruitment to mRNA through this canonical initiation pathway.

Our studies of the dynamics of ribosome interactions with *hp-capCAA*_*AUG*_ mRNAs provide insight into the modulation of cap-dependent initiation by m^7^G cap-distal secondary structures in 5′ UTR. Earlier work showed that these hairpins inhibit translation in a stability-dependent manner (10, 12). We observed a similar correlation with 60S-mRNA binding dynamics, which showed reduced 60S recruitment with increased hairpin stability (Figures 3*D* and 3*E*). While these results are consistent with the ability of stable m^7^G cap-distal hairpins to inhibit translation by impeding 43S scanning, the mechanism of such inhibitory effect was not known. Intriguingly, we observed two modes of 40S-mRNA interactions on *hp-capCAA*_*AUG*_ mRNAs. 40S binding events through one mode were short-lived (dwell time < 100 sec) and non-productive for 60S recruitment, whereas the other mode showed long-lived 40S binding events (dwell time > 100 sec) that were productive for 60S recruitment (Figures 4C and 5). Stable hairpins promote the non-productive short-lived mode of 40S-mRNA interaction and suppress the productive long-lived mode of 40S-mRNA interaction (Figures 4C and 5). Collectively, our results indicate that stable m^7^G cap-distal hairpins reduce 60S recruitment by promoting premature 43S dissociation from mRNA.

The long-lived (>100 sec) 40S binding with CAA-repeat mRNA (Figure 4C) and the predominance of single 40S binding to these mRNAs (Figure 6) support a cap-tethered scanning model for initiation (23), which postulates that 43S/eIF4F interactions with 5’ cap are maintained until initiation has been completed. This model also provides an explanation for the ability of 8–16 GC base pair hairpins to increase the numbers of 40S binding events per trajectory (Figures 4A,B). Since the 8–16 GC base pair hairpins induce premature 40S dissociation (Figure 4D), stable hairpins may disrupt 43S/eIF4F/5’ cap interactions and thereby promote 5’ cap availability for additional 43S recruitment.

To generate the cell-free translation system with reconstituted extract and fluorescently labeled ribosomes, we employed extract fractionation and maleimide dye ribosome-labeling techniques. Since these techniques do not require specialized translation systems or genetically modified ribosomes (25, 26), our approach should provide a general method for generating *in vitro* translation systems with fluorescently labeled ribosomal subunits. In addition to hairpin-mediated effects on ribosome dynamics, the fluorescent ribosome translation system can be applied to study the modulation of ribosome dynamics by other 5′ UTR-localized elements, such as more complex secondary structures (2, 5, 27), upstream open reading frames (28), inhibitory codon pairs (29), and RNA-binding proteins (30). This method for generating a fluorescent ribosome translation system should enable broad applications for studying ribosome recruitment dynamics during cap-dependent initiation from diverse eukaryotic systems.

## MATERIALS AND METHODS

### Preparation of reconstituted translation extract with fluorescently labeled ribosomes

Translation extracts derived from *S. cerevisiae* strain YAS1874 (*MAT***a** *MAK10::URA3 PEP4::HIS3 prb1 prc1 ade2 trp1 his3 ura3*) (31) were prepared as previously described (32). Triplet-state quencher-conjugated maleimide dyes (14) were used to label high-salt washed 80S ribosomes and the fluorescently labeled 40S and 60S subunits were purified from high-salt sucrose gradients (33). Fluorescently labeled 40S (LD550-40S) and 60S (LD650-60S) were combined with ribosome depleted extract and extract factors to reconstitute the extract with fluorescently labeled ribosomal subunits.

#### i) Labeling and purification of ribosomal subunits

Extract (430 μl) was diluted with an equal volume of buffer B (30 mM HEPES-KOH, pH 7.4; 15 mM MgOAc, pH 6; 100 mM KOAc, pH 8, 0.1 mM PMSF), layered on a high-salt sucrose cushion (buffer B with 500 mM KCl, 1 mg/ml heparin, 1 M sucrose), and ribosomes were pelleted by ultracentrifugation with a TLA100.2 rotor for 2 h at 90,000 rpm and 4 °C. At 4 °C, the pellet was rinsed twice, each with 300 μl of buffer B, and then suspended with 300 μl of buffer B. The ribosome suspension was transferred to a microfuge tube, centrifuged for 30 sec at 14,000 rpm and 4 °C to pellet debris, and the supernatant was collected. Absorbance at 260 nm was measured and an extinction coefficient of 5 × 10^7^ cm^-1^ M^-1^ was used to determine the 80S ribosome concentration. Ribosomes were maintained on ice unless indicated otherwise.

The ribosomes (∼1.2 nmol) at a concentration of 0.7 μM were fluorescently labeled in 140 μM of either LD550-MAL or LD650-MAL (Lumidyne Technologies) for 1 h on ice followed by addition of 1.4 mM 2-mercaptoethanol. The ribosome suspension buffer was then diluted 70-fold with buffer B, 35-fold with buffer A (30 mM HEPES-KOH, pH 7.4; 3 mM MgOAc, pH 6; 100 mM KOAc, pH 8, 0.1 mM PMSF; 2 mM DTT), and 35-fold with subunit separation buffer (50 mM HEPES-KOH, pH 7.4; 2 mM MgCl_2_; 500 mM KCl; 0.1 mM PMSF; 2mM DTT) using 100 kD Amicon Ultra-4 filters pre-equilibrated with buffer B. The A_260_ was measured and the ribosomes at a concentration of ∼0.8 μM were incubated in 1 mM puromycin on ice for 15 min then at 37 °C for 10 min. The ribosomes were then layered onto a 10–30% high-salt sucrose gradient (50 mM HEPES-KOH, pH 7.4; 5 mM MgCl_2_; 500 mM KCl; 0.1 mM EDTA; 10% or 30% sucrose, 2mM DTT), centrifuged at 22,000 rpm for 17 h at 4 °C in a Surespin-630 rotor, and fractionated while scanned at A_254_. Fractions containing 40S and 60S were collected and 100K MWCO Amicon Ultra-4 filters (Millipore) were used to exchange the sucrose gradient with buffer A until the KCl concentration was ∼10 nM. The 40S and 60S concentrations were determined by measuring the A_260_ and using the extinction coefficients of 2 × 10^7^ cm^-1^ M^-1^ and 4 × 10^7^ cm^-1^ M^-1^ for 40S and 60S subunits, respectively. The subunits were aliquoted, frozen in liquid N_2_, and stored at –80 °C.

#### ii) Preparation of Extract Factors

A 1.0 ml frozen aliquot of extract was thawed on ice, concentrated to ∼60 μl with a 100K MWCO Amicon Ultra-0.5 filter (Millipore) pre-equilibrated with buffer A, eluted, and mixed with 340 μl of buffer C (30 mM HEPES-KOH, pH 7.4; 3 mM MgOAc, pH 6; 100 mM KOAc, pH 8; 590 mM KCl; 0.1 mM PMSF; 2 mM DTT). The extract (∼400 μl) was then layered onto a 10–30% high-salt sucrose gradient and centrifuged as described above. The top ∼15 ml of the gradient was collected, concentrated to ∼1.0 ml in a 3K MWCO Amicon Ultra-15 filter (Milllipore) pre-equilibrated with buffer A, transferred to a 3K MWCO Amicon Ultra-0.5 filter (Milllipore) pre-equilibrated with buffer A, and the buffer was then exchanged by diluting the concentrate with buffer A supplemented with 5% glycerol until the KCl concentration was ∼1.0 μM. The final volume of extract factors was adjusted to 100 μl with buffer A supplemented with 5% glycerol. Extract factors were then aliquoted, frozen in liquid N_2_, and stored at –80 °C.

#### iii) Preparation of Ribosome-Depleted Extract

Ribosome-depleted extract (RDE) was generated from 1.2 ml of extract that was incubated with 0.5 U/ml of micrococcal nuclease (Sigma) at 25 °C for 5 min, as described previously (34). Nuclease-treated extract was used because it produced RDE that increased the absolute level of reporter RNA translation in reconstituted bulk translation reactions. Ribosomes in the nuclease-treated extract were pelleted by ultracentrifugation with a TLA100.2 rotor at 90,000 rpm for 1 h at 4 °C. The top 1.1 ml of the supernatant was transferred to a new chilled ultracentrifuge tube and residual ribosomes remaining in the supernatant were pelleted by another round of ultracentrifugation at 90,000 rpm for 1 h at 4 °C. The top 1.0 ml of the supernatant was aliquoted, frozen with liquid N_2_, and stored at –80°C.

#### iv) Extract reconstitution and translation mix assembly

To reconstitute *S. cerevisiae* extract with fluorescently labeled ribosomes, 3.8 μl of buffer A supplemented with 5% glycerol was combined with 0.2 pmol LD550-40S, 0.2 pmol LD650-60S, 0.4 μl extract factors, and 9 μl RDE to give a final reconstituted extract volume of 14 μl. The reconstituted extract was prepared as single-use aliquots, frozen in liquid N_2_, and stored at –80°C. 20 μl of translation mix was assembled fresh by combining 14 μl reconstituted extract with 6 μl of master mix (32). The master mix provides the translation mix and final translation reaction with 2.9 mM and 170 mM of Mg^2+^ and K^+^ glutamate salts, respectively.

### RNA synthesis and analysis

All mRNAs were synthesized with the MEGAscript T7 Kit (Ambion) according to the manufacturer’s protocol. DNA templates for synthesis of *LUC* mRNAs were prepared as described previously (32). Synthesis of *capCAA*_*AUG*_, *capCAA*_*AAC*_, and *hp-capCAA*_*AAC*_ mRNAs was performed with synthetic DNA templates (IDT) and a clamp DNA oligo (IDT) that was complimentary to the 3′ end of each CAA repeat DNA template and contained a T7 RNA polymerase binding sequence (Supplementary Table 2). 5′-end capping and 3′-end biotinylation were performed using the Vaccinia Capping System (New England Biolabs) and the Pierce RNA 3′ End Biotinylation Kit (Thermo Scientific), respectively. All mRNAs were phenol-chloroform extracted and subsequently purified with the Direct-Zol RNA kit (Zymo Research), from which mRNA was eluted in water. Integrity and concentration of mRNA were assessed by denaturing (8M urea) 5% acrylamide gel electrophoresis and SYBR green II RNA staining.

### Single-molecule experiments

Assembly of single-molecule detection chambers derivatized with mPEG, biotin-PEG, and MS-(PEG)_4_ has been described previously (35, 36). Individual flow channels were incubated with 10 μl of 0.2 μg/μl streptavidin (Thermo Scientific) for 10 min followed by three washes, each with 15 μl of T50 buffer (20 mM Tris-HCl (pH 7.0), 50 mM NaCl). Streptavidin-treated channels were flushed with 20 μl of 3′-end biotinylated mRNA diluted in T50 buffer to 1 – 3 ng/μl, incubated for 15 min, and flushed three times, each with 15 μl of T50 buffer. Channels were then flushed with two sequential 15 μl additions of 0.5 nM Cy3-CAA oligo (5′-CAACAACAACAACAACAA-3′) and incubated for 5 min to fluorescently label immobilized mRNA. Unbound Cy3-CAA oligo was removed by three 15 μl washes with T50 buffer and fluorescence was imaged to assess relative mRNA density in channels. All flow channel incubations were performed at 25 °C. Syringe pump delivery of translation reaction mixtures to flow channels and objective-type TIRF imaging conditions were described previously (35, 36). The 532 and 640 nm laser illumination intensities were 10 and 4 μW, respectively. Data were recorded as a kinetic series at a speed of 2 seconds per frame. Image analyses were performed using custom written Matlab codes, as described previously (32, 36).

## ACKNOWLEDGMENTS

This work was supported by the National Institutes of Health [R01GM121847], the Memorial Sloan Kettering Cancer Center (MSKCC) Support Grant/Core Grant [P30 CA008748], and the MSKCC Functional Genomics Initiative.

## REFERENCES

1. Merrick WC & Pavitt GD (2018) Protein Synthesis Initiation in Eukaryotic Cells. Cold Spring Harb Perspect Biol 10(12).

2. Fay MM, Lyons SM, & Ivanov P (2017) RNA G-Quadruplexes in Biology: Principles and Molecular Mechanisms. J Mol Biol 429(14):2127–2147.

3. Svoboda P & Di Cara A (2006) Hairpin RNA: a secondary structure of primary importance. Cell. Mol. Life Sci. 63(7-8):901–908.

4. Tan D, Marzluff WF, Dominski Z, & Tong L (2013) Structure of histone mRNA stem-loop, human stem-loop binding protein, and 3’hExo ternary complex. Science 339(6117):318–321.

5. Varshney D, Spiegel J, Zyner K, Tannahill D, & Balasubramanian S (2020) The regulation and functions of DNA and RNA G-quadruplexes. Nat Rev Mol Cell Biol 21(8):459–474.

6. Jodoin R, Carrier JC, Rivard N, Bisaillon M, & Perreault JP (2019) G-quadruplex located in the 5’UTR of the BAG-1 mRNA affects both its cap-dependent and cap-independent translation through global secondary structure maintenance. Nucleic Acids Res 47(19):10247–10266.

7. Lee AS, Kranzusch PJ, & Cate JH (2015) eIF3 targets cell-proliferation messenger RNAs for translational activation or repression. Nature 522(7554):111–114.

8. Sabarinathan R, et al. (2014) Transcriptome-wide analysis of UTRs in non-small cell lung cancer reveals cancer-related genes with SNV-induced changes on RNA secondary structure and miRNA target sites. PLoS One 9(1):e82699.

9. Wolfe AL, et al. (2014) RNA G-quadruplexes cause eIF4A-dependent oncogene translation in cancer. Nature 513(7516):65–70.

10. Kozak M (1989) Circumstances and mechanisms of inhibition of translation by secondary structure in eucaryotic mRNAs. Mol Cell Biol 9(11):5134–5142.

11. Babendure JR, Babendure JL, Ding JH, & Tsien RY (2006) Control of mammalian translation by mRNA structure near caps. RNA 12(5):851–861.

12. Kozak M (1986) Influences of mRNA secondary structure on initiation by eukaryotic ribosomes. Proc Natl Acad Sci U S A 83(9):2850–2854.

13. Mitchell SF, et al. (2010) The 5’-7-methylguanosine cap on eukaryotic mRNAs serves both to stimulate canonical translation initiation and to block an alternative pathway. Mol Cell 39(6):950–962.

14. Zheng Q, et al. (2014) Ultra-stable organic fluorophores for single-molecule research. Chem. Soc. Rev. 43(4):1044–1056.

15. Tarun SZ, Jr. & Sachs AB (1995) A common function for mRNA 5’ and 3’ ends in translation initiation in yeast. Genes Dev 9(23):2997–3007.

16. Brar GA, et al. (2012) High-resolution view of the yeast meiotic program revealed by ribosome profiling. Science 335(6068):552–557.

17. Ingolia NT, Ghaemmaghami S, Newman JR, & Weissman JS (2009) Genome-wide analysis in vivo of translation with nucleotide resolution using ribosome profiling. Science 324(5924):218–223.

18. Ingolia NT, Lareau LF, & Weissman JS (2011) Ribosome profiling of mouse embryonic stem cells reveals the complexity and dynamics of mammalian proteomes. Cell 147(4):789–802.

19. Sobczak K, et al. (2010) Structural diversity of triplet repeat RNAs. J Biol Chem 285(17):12755–12764.

20. Zuker M (2003) Mfold web server for nucleic acid folding and hybridization prediction. Nucleic Acids Res 31(13):3406–3415.

21. Yourik P, et al. (2017) Yeast eIF4A enhances recruitment of mRNAs regardless of their structural complexity. Elife 6.

22. Pelletier J & Sonenberg N (1985) Insertion mutagenesis to increase secondary structure within the 5’ noncoding region of a eukaryotic mRNA reduces translational efficiency. Cell 40(3):515–526.

23. Archer SK, Shirokikh NE, Beilharz TH, & Preiss T (2016) Dynamics of ribosome scanning and recycling revealed by translation complex profiling. Nature 535(7613):570–574.

24. Wu CC, Zinshteyn B, Wehner KA, & Green R (2019) High-Resolution Ribosome Profiling Defines Discrete Ribosome Elongation States and Translational Regulation during Cellular Stress. Mol Cell 73(5):959–970 e955.

25. Jakubowicz T, Palen E, & Gasior E (1978) The effect of sulfhydryl reagents on the activity and stability of yeast ribosomes. Acta Biochim. Pol. 25(1):49–59.

26. Lee T & Heintz RL (1976) Reaction of N-(3-pyrene)maleimide with thiol groups of reticulocyte ribosomes. Eur J Biochem 66(1):105–114.

27. Guenther UP, et al. (2018) The helicase Ded1p controls use of near-cognate translation initiation codons in 5’ UTRs. Nature 559(7712):130–134.

28. Barbosa C, Peixeiro I, & Romao L (2013) Gene expression regulation by upstream open reading frames and human disease. PLoS Genet 9(8):e1003529.

29. Tesina P, et al. (2020) Molecular mechanism of translational stalling by inhibitory codon combinations and poly(A) tracts. EMBO J 39(3):e103365.

30. Medenbach J, Seiler M, & Hentze MW (2011) Translational control via protein-regulated upstream open reading frames. Cell 145(6):902–913.

31. Kessler SH & Sachs AB (1998) RNA recognition motif 2 of yeast Pab1p is required for its functional interaction with eukaryotic translation initiation factor 4G. Mol Cell Biol 18(1):51–57.

32. Gaba A, Wang H, Fortune T, & Qu X (2020) Smart-ORF: a single-molecule method for accessing ribosome dynamics in both upstream and main open reading frames. Nucleic Acids Res.

33. Acker MG, Kolitz SE, Mitchell SF, Nanda JS, & Lorsch JR (2007) Reconstitution of yeast translation initiation. Methods Enzymol 430:111–145.

34. Wu C & Sachs MS (2014) Preparation of a Saccharomyces cerevisiae cell-free extract for in vitro translation. Methods Enzymol. 539:17–28.

35. Wang H, Sun L, Gaba A, & Qu X (2020) An in vitro single-molecule assay for eukaryotic cap-dependent translation initiation kinetics. Nucleic Acids Res 48(1):e6.

36. Gaba A, Wang HY, & Qu XH (2020) An In Vitro Single-Molecule Imaging Assay for the Analysis of Cap-Dependent Translation Kinetics. Jove-J Vis Exp (163).

