## Supplementary Material for "Stable m^7^G Cap-Distal 5′UTR Hairpin Structure Mediates Distinct 40S and 60S Binding Dynamics"

**SUPPLEMENTARY FIGURE 1**

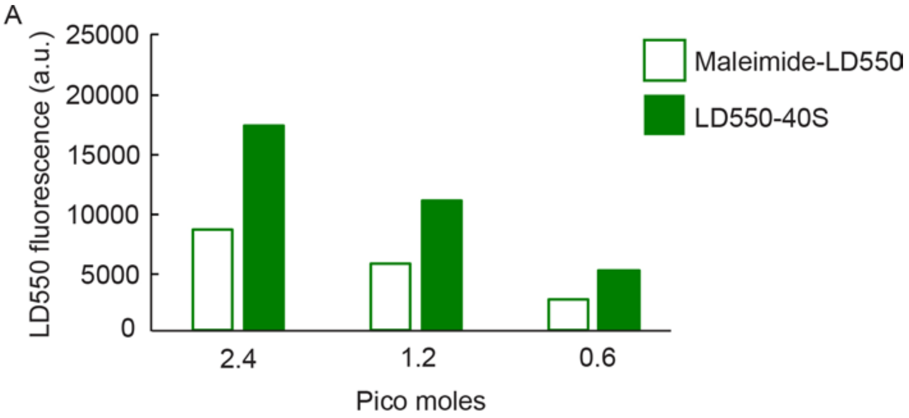


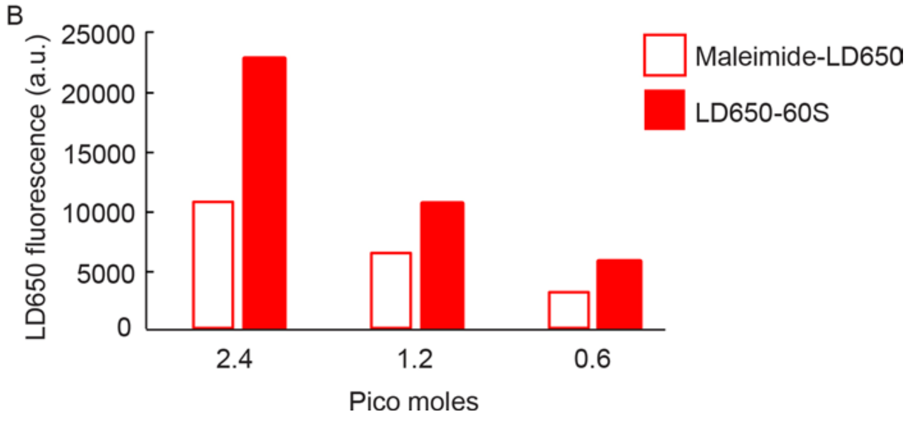


**Supplementary Figure 1. Fluorescence levels from LD550- and LD650-maleimide dyes and LD550-40S and LD650-60S ribosomal subunits.** (*A* and *B*) The fluorescence from 2.4, 1.2, and 0.6 pmols of each maleimide dye and of each fluorescently labeled ribosomal subunit was quantified by densitometry. Fluorescence levels from equal molar amounts of LD550 and LD550-40S (A) and LD650 and LD650-60S (B) are plotted as indicated.

### Supplementary Table 1

**Table 1. Thermostability of GC base pair hairpins**

| Hairpin stem (base pairs) | Sequence* | Minimum free energy |
|---|---|---|
| 4 GC | CCGGACACCCGG | -5.4 kcal/mol |
| 8 GC | CGCCGGCGACACCGCCGGCG | -16.5 kcal/mol |
| 12 GC | CGCCCGCCGGCGACACCGCCGGCGGGCG | -28.9 kcal/mol |
| 16 GC | CGGCCGCCCGCCGGCGACACCGCCGGCGGGCGGCCG | -41.3 kcal/mol |

*The hairpin loop sequence is underlined.

### Supplementary Table 2

| | DNA template sequence* |
|---|---|
| $CAA_{AUG}$ ( 0 GC $hp\text{-}CAA_{AUG}$ ) | 5'-TTGTTGTTGTTGTTGTTGTTGTTGTTGTTGTTGTTGTTGTTGTTGTTGTTGTTG TTGTTCCATTTTGTTGTTGTTGTTGTTGTTGTTGTTGTTGTTGTTGTTGTTGTTGT TGTTGTTGTTGTT**CCTATAGTGAGTCGTATTACATATGCGTGTTACC** -3' |
| $CAA_{AAC}$ | 5'-GTTGTTGTTGTTGTTGTTGTTGTTGTTGTTGTTGTTGTTGTTGTTGTTGTTGTT GTTGTTGTTGTTGTTGTTGTTGTTGTTGTTGTTGTTGTTGTTGTTGTTGTTGTTGT TGTTGTTGTTGTT**CCTATAGTGAGTCGTATTACATATGCGTGTTACC**-3' |
| 4 GC $hp\text{-}CAA_{AUG}$ | 5'-TTGTTGTTGTTGTTGTTGTTGTTGTTGTTGTTGTTGTTGTTGTTGTTGTTGTTG TTGTTCCATTTTGTTGTTGTTCCGGGTGTCCGGGTTGTTGTTGTTGTTGTTG TTGTTGTTGTTGTTGTTGTTGTTGTT**CCTATAGTGAGTCGTATTACATATGC GTGTTACC**-3' |
| 8 GC $hp\text{-}CAA_{AUG}$ | 5'-TTGTTGTTGTTGTTGTTGTTGTTGTTGTTGTTGTTGTTGTTGTTGTTGTTGTTG TTGTTCCATTTTGTTGTTGTTCGCCGGCGGTGTCGCCGGCGGTTGTTGTTG TTGTTGTTGTTGTTGTTGTTGTTGTTGTTGTTGTT**CCTATAGTGAGTCGTAT TACATATGCGTGTTACC**-3' |
| 12 GC $hp\text{-}CAA_{AUG}$ | 5'-TTGTTGTTGTTGTTGTTGTTGTTGTTGTTGTTGTTGTTGTTGTTGTTGTTGTTG TTGTTCCATTTTGTTGTTGTTCGCCCGCCGGCGGTGTCGCCGGCGGGCGG TTGTTGTTGTTGTTGTTGTTGTTGTTGTTGTTGTTGTTGTTGTTGTT**CCTATAGT GAGTCGTATTACATATGCGTGTTACC**-3' |
| 16 GC $hp\text{-}CAA_{AUG}$ | 5'-TTGTTGTTGTTGTTGTTGTTGTTGTTGTTGTTGTTGTTGTTGTTGTTGTTGTTG TTGTTCCATTTTGTTGTTGTTCGGCCGCCCGCCGGCGGTGTCGCCGGCGG GCGGCCGGTTGTTGTTGTTGTTGTTGTTGTTGTTGTTGTTGTTGTTGTTGTTGTT **CCTATAGTGAGTCGTATTACATATGCGTGTTACC**-3' |
| DNA clamp | 5'-GGTAACACGCATATGTAATACGACTCACTATAGG-3' |

*The sequences for the AUG start site and AUG → AAC mutation are shown in red. The sequence complimentary to the clamp oligo is in bold.
